# Young APPKI ^NL-G-F/NL-G-F^ mice display high-fat diet-induced metabolic disturbances and specific disorders associated with brain energy homeostasis

**DOI:** 10.1101/2021.12.21.473697

**Authors:** Wei Wang, Daisuke Tanokashira, Megumi Maruyama, Chiemi Kuroiwa, Takashi Saito, Takaomi C. Saido, Akiko Taguchi

## Abstract

**Aim:** Type 2 diabetes mellitus (T2DM) is an increased risk factor for Alzheimer’s disease (AD); however, the relationship between the two conditions is controversial. High-fat diet (HFD) causes cognitive impairment with/without Aβ accumulation in middle-aged or aged transgenic (Tg) and knock-in (KI) AD mouse models, except for metabolic disorders, which commonly occur in all mice types. Alternatively, whether HFD in early life impacts energy metabolism and neurological phenotypes in young AD mouse models remains unknown. In the present study, we examined the effects of HFD on young APPKI ^NL-G-F/NL-G-F^ mice, one of the novel knock-in (KI)-AD mouse models.

**Methods:** The mice were categorized by diet into two experimental groups, normal diet (ND) and HFD. Four-week-old WT and APPKI ^NL-G-F/NL-G-F^ mice were fed ND or HFD for nine weeks. Both types of mice on ND and HFD were examined during young adulthood.

**Results:** HFD causes T2DM-related metabolic disturbances in young WT and APPKI ^NL-G-F/NL-G-F^ mice and specific impairment of brain energy homeostasis only in young APPKI ^NL-G-F/NL-G-F^ mice. However, HFD-induced metabolic dysfunctions had no impact on behaviors, Aβ levels, and specific IRS1 modifications in both young APPKI ^NL-G-F/NL-G-F^ mice and young WT mice.

**Conclusion:** HFD in early life is effective in causing metabolic disturbances in young WT and APPKI ^NL-G-F/NL-G-F^ mice but is ineffective in inducing neurological disorders in young mice, which suggests that the aging effects along with long-term HFD cause neurological alterations.

## 1 INTRODUCTION

Epidemiological evidence and experimental studies on animal models have suggested that type 2 diabetes mellitus (T2DM), characterized by impaired glucose metabolism and insulin resistance, is one of the risk factors for dementia, including Alzheimer’s disease (AD)^1, 2^. Aberrant alterations of the insulin signaling pathway mediated by insulin receptor substrate proteins correlate with the onset of T2DM ^3^. Recently, the IRS1 phosphorylation at specific serine (Ser) sites, such as human(h)Ser312/mouse(m)Ser307 implicated in metabolic dysfunctions and human(h)Ser616/mouse(m)Ser612, human(h)Ser636/mouse(m)Ser632, and human(h)Ser639/mouse(m)Ser635 known as negative regulatory sites for IRS1 signaling has been found in postmortem AD brain tissues^4, 5^. Additionally, our previous studies have shown that these modifications of neural IRS1 occur in the brains of young amyloid precursor protein (APP) knock-in (KI) ^NL-G-F/NL-G-F^ (APPKI ^NL-G-F/NL-G-F^) mouse model. This novel mouse model of AD uses a KI strategy, which overcomes the artificial effects of transgenes of transgenic (Tg) AD mouse models carrying humanized APP with Swedish NL, Beyreuther/Iberian F, and Arctic mutations. The mouse model also exhibits normal energy metabolism, normal cognitive functions, and increased Aβ levels from 2 months of age, and cognitive deficits after six months^6, 7^. These results suggest that specific IRS1 modifications are involved in the original elevation of Aβ42 level but not cognitive deficits in these mice^7^.

Alternatively, previous clinical research has revealed a controversial relationship between T2DM and AD ^8-10^. Similarly, basic research on conventional AD mouse models using transgenic (Tg) strategies have shown that a high-fat diet (HFD) that mimics the physiological status of T2DM, leads to cognitive impairment, regardless of the extent of the disorder, with/without alteration of Aβ accumulation in Tg-AD mouse models, such as Tg2576 and 3xTg-ADmice^11-13^. Moreover, in middle-aged APPKI ^NL/N^ mouse model with Swedish NL mutation, a long-term HFD (60% calories from fat) from 2 to 18 months of age leads to mild behavioral impairment without Aβ production and exhibits normal cognitive function in late middle ages without Aβ deposition^14^. sChronic HFD (40% calories from fat) from 6 to 12 months of age promotes Aβ aggregation and cognitive decline in middle-aged APPKI ^NL-F/NL-F^ mice, carrying humanized APP with Swedish NL and Beyreuther/Iberian F mutations at mouse APP locus. This model displays Aβ accumulation from 6 months of age and very mild cognitive deficits at 18 months^15^. However, metabolic disorders are commonly observed in both Tg and KI AD mouse models on HFD^6, 11-16^.

Nevertheless, whether exposure to hyperalimentation in the early life of APPKI mice impacts the asymptomatic state in energy metabolism and cognitive function in addition to Aβ levels and neural IRS1 modifications remains unknown. To address this question, we have examined the effects of HFD (60% calories from fat) on young APPKI ^NL-G-F/NL-G-F^ mice. We found that HFD for nine weeks leads to hyperglycemia and hyperlipidemia in young WT and APPKI ^NL-G-F/NL-G-F^ mice with impairment of ketogenesis only in young APPKI ^NL-G-F/NL-G-F^ mice. However, HFD-induced metabolic disorders do not impair cognitive functions, alter the Aβ or phosphorylation levels of hippocampal IRS1 in both young WT and APPKI ^NL-G-F/NL-G-F^ mice. These results suggest that HFD in early life is effective in causing metabolic cgical manifestations in young APPKI ^NL-G-F/NL-G-F^ mice and young WT mice.

## 2 METHODS

### 2.1 Animals

C57BL/6J male wild-type (WT) mice supplied by Japan SLC, Inc. (Shizuoka, Japan) were used to establish type 2 diabetes mellitus (T2DM) mice and their respective control mice. APP KI^NL-G-F/NL-G-F^ (Swedish (NL), Arctic (G), and Beyreuther/Iberian (F) mutations) homozygous mice were obtained from Dr. Saido at the Laboratory for Proteolytic Neuroscience, RIKEN Brain Science Institute, Saitama, Japan^6^. The WT and APP KI^NL-G-F/NL-G-F^ mice were categorized into two experimental groups: normal diet (ND) and high-fat diet (HFD) groups. The ND group was fed with ND (CE-2; CLEA Japan Inc., Tokyo, Japan) and HFD group was fed with HFD (D12492, 60% kcal from fat; Research Diets, Inc., New Brunswick, NJ, USA) for 9 weeks, respectively. All the mice were housed at room temperature (25°C ± 2°C) under a standard 12/12 h light– dark cycle with free access to water and food. Animal experiments were conducted in compliance with the guidelines and approval of the ethics committee in Animal Care and Use of the National Center for Geriatrics and Gerontology in Japan.

### 2.2 Measurement of metabolic parameters

Body weight and blood glucose were measured weekly throughout the study. Blood glucose was measured using a portable glucose meter (ACCU-CHEK® Aviva, Roche DC Japan K.K., Tokyo, Japan). The plasma level of insulin at 6 h fasting was determined using an insulin enzyme-linked immunosorbent assay (ELISA) kit (Morinaga, Yokohama, Japan). During random feeding and 3 h fasting, rectal temperature was measured using a rectal probe (Animal warmer, thermometer, Bio Research Center Co., LTD, Japan). The daily food intake was recorded after a 24 h fast. The biochemical parameters, including total cholesterol, low-density lipoprotein (LDL)-cholesterol, high-density lipoprotein (HDL)-cholesterol, free fatty acid, total ketone bodies, and triglyceride in 6 h fasting plasma, were measured using enzymatic methods (Oriental Yeast Co., Ltd., Japan).

### 2.3 Glucose tolerance tests and insulin tolerance tests

During the glucose tolerance tests, the mice were fasted overnight for 24 h and injected intraperitoneally with glucose at 1.5 g/kg body weight (Otsuka, Japan). Blood glucose was measured at 0, 15, 30, 60, and 120 min after injection. During the insulin tolerance tests, the mice were fasted briefly for 3 h and injected intraperitoneally with insulin at 0.75 units/kg body weight (Humulin R, Eli Lilly Japan K.K.). Blood glucose was collected at 0, 30, 60, 90, and 120 min after injection and normalized to the initial value at 0 min.

### 2.4 Behavior

#### 2.4.1 Open field

The mice were gently placed in the center of a circular open field having a diameter of 100 cm, and allowed to move freely for 5 min. The total distance a mouse traveled was recorded and analyzed via a video camera connected to the system software (O’HARA & CO., LTD, Japan).

#### 2.4.2 Elevated plus maze

The elevated plus maze consists of two 25 × 5 cm open arms opposite to two 25 × 5 cm closed arms enclosed by a 20 cm wall. The arms extended from a 5 × 5 cm central platform and the maze was had an elevation of50 cm above the floor. The mice were first placed on the central platform of the maze and allowed to move freely for 5 min. Their behaviors were continuously recorded with a video and then analyzed with the system software (O’HARA & CO., LTD, Japan). The time spent by the mice in the open arms was defined as anxiety-related behaviors.

#### 2.4.3 Passive avoidance

Passive avoidance was performed using a step-through box (Harvard Apparatus, USA) consisting of a lighted chamber (20 × 20 × 25 cm) and a dark chamber (7 × 15 × 10 cm) connected with a sliding door. At first, mice were placed in the lighted compartment. sWhen the mice entered the dark chamber, the sliding door was closed, and they received a light shock at the foot (0.15 mA, 1 s). Mice were trained up to criterion defined as the mice remaining in the lighted compartment for 120 s. The number of trials needed to reach this criterion was taken as the measure of task acquisition.

#### 2.4.4 Water T maze

Hippocampal-dependent spatial memory of the mice was tested using a water T maze according to our previous reports^7, 17, 18^. For the first screening step, the preference of each mouse was determined. The mice were placed in the start box and allowed to swim to the right or left arm. This screening step was repeated three times at 30 s intervals. The platform was placed on the side that the mice reached less often. Next, the mice were allowed to explore the right and left sides of the maze freely. If mice reached the platform (correct choice), they rested for 5 s on the platform. If not, the arm entry was closed with a board (incorrect choice), and they received a deterrent by being forced to swim for 15 s. This trial step was repeated five times at 5 min intervals. The mice were subjected to the trial step for five days. The percentage of correct responses per day was determined.

### 2.5 Western blotting

Western blot analysis was conducted as previously described protocol^7^. In summary, hippocampal tissue was homogenized in a lysis buffer T-PER^®^ Tissue Protein Extraction Reagent (Thermo Fisher Scientific, Waltham, MA, USA) containing a protease inhibitor cocktail (Nacalai Tesque, Kyoto, Japan) and a phosphatase inhibitor cocktail (Nacalai Tesque) with a pellet mixer. After incubation on ice for 15 min, the lysates were centrifuged for 5 min at 14,200 × g and 4°C. Protein concentrations were determined using a BCA protein assay kit (Pierce, Rockford, IL, USA). The protein samples were separated by SDS-PAGE and then transferred to a polyvinylidene fluoride membrane. sThe membranes were blocked using Block Ace (Yukijirushi Ltd., Sapporo, Japan) and incubated with the indicated primary antibodies: rabbit anti-phospho-IRS1 [mouse Ser307/human Ser312 (mSer307/hSer312), mouse Ser612/human Ser616 (mSer612/hSer616), mouse Ser632/Ser635/human Ser636/Ser639 (mSer632/Ser635/hSer636/Ser639), mouse Ser1097/human Ser1101 (mSer1097/hSer1101)] {1:400; Cell Signaling Technology(CST)}, rabbit anti-IRS1 (1:1000; CST). Immunodetection was performed with horseradish peroxidase-conjugated secondary antibodies (1:5000; CST).The chemiluminescent signals were detected using Chemi-Lumi One (Nacalai Tesque) or ImmunoStar LD (FUJIFILM Wako). The images were scanned using Amersham Imager 680 (GE Healthcare Life Sciences, UK).

### 2.6 ELISA quantitation of Aβ

Quantitation of Aβ levels was conducted as previously described^7^. In summary, the levels of T-PER-extractable Aβ40 and Aβ42 in the hippocampus tissues were determined using the Human/Rat/Mouse β Amyloid (1–40) ELISA Kit Wako II (#294-64701; Fujifilm Wako Pure Chemical Corp., Osaka, Japan) and the Human/Rat/Mouse β Amyloid (1–42) ELISA Kit Wako, High Sensitivity (#292-64501, Fujifilm Wako Pure Chemical Corp.) as per the manufacturers’ instructions.

### 2.7 Statistical analysis

All results were expressed as mean ± standard error of the mean (SEM) in the text. Statistical analyses were performed using Prism7 for Mac OS X v.7.0d (GraphPad Software Inc., USA). Comparisons of means from multiple groups were analyzed using one-way analysis of variance (ANOVA) or two-way ANOVA followed by Tukey– Kramer’s test and Bonferroni’s test. Significance was indicated as * p < 0.05, ** p < 0.01; †p < 0.05, ††p < 0.01; ‡ p < 0.05, ‡‡ p < 0.01; § p < 0.05, §§ p < 0.01; ¶ p < 0.05, ¶¶ p< 0.01.

## 3 RESULTS

### 3.1 HFD leads to metabolic disturbance in both young WT and APPKI ^NL-G-F/NL-G-F^ mice

To examine the effects of HFD on young APPKI ^NL-G-F/NL-G-F^ mice from an early age to young adulthood, 4-week-old WT and APPKI ^NL-G-F/NL-G-F^ mice were fed a ND (12% calories from fat) or HFD (60% calories from fat) for nine weeks. At baseline, there were no differences in body weight or random blood glucose levels between both mice on ND (Figure 1A and B), which was consistent with the data from our previous studies ^7^. Compared with ND groups, both mice on HFD showed a similar increase in body weight despite decreased food intake (Figure 1A and F). Although insulin sensitivity reduced only in HFD-WT mice (Figure 1D), HFD-APPKI ^NL-G-F/NL-G-F^ mice displayed a more severe glucose intolerance than HFD-WT mice, while both WT and APPKI ^NL-G-F/NL-G-F^ mice on ND had normal glucose tolerance (Figure 1C). Consistent with these results, normal levels of blood insulin and hyperinsulinemia were observed in the ND group and in the HFD group, respectively (Figure 1E). Interestingly, ND-APPKI ^NL-G-F/NL-G-F^ mice exhibited a significant decrease in core body temperature correlated with energy metabolism; however, it was no longer observed under HFD and/or 3h fasting conditions (Figure 1G). Also, compared with young WT mice, we found that young APPKI ^NL-G-F/NL-G-F^ mice displayed a consistent reduction in the levels of fasting total blood ketone bodies, which in general rise under fasting or T2DM regardless of diet type (Figure 1H). In contrast, increased levels of total, low-density lipoprotein (LDL), and high-density lipoprotein (HDL) blood cholesterol decreased levels of blood triglyceride were observed (corresponding to decreased food intake in both mice on HFD), while the levels of plasma free fatty acid were comparable between all groups (Figure 1I, J, K). These results indicate that HFD in early life causes T2DM-related typical metabolic disorders in both young WT and APPKI ^NL-G-F/NL-G-F^ mice and disorders specific to energy homeostasis in young APPKI ^NL-G-F/NL-G-F^ mice.

**FIGURE 1.**
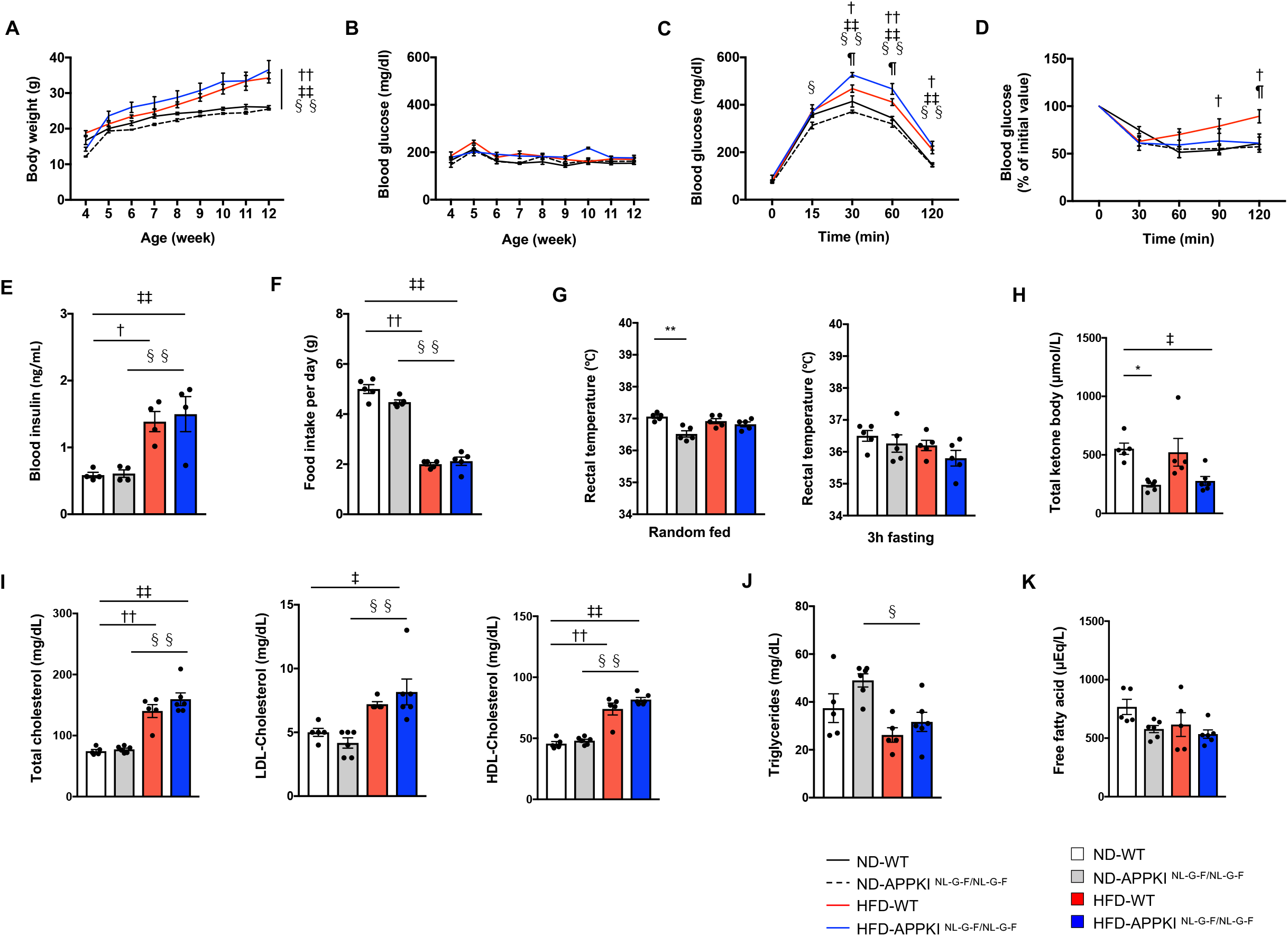
Changes in metabolic parameters in high-fat diet (HFD)-induced young APPKI ^NL-G-F/NL-G-F^ mice. (A) Body weight and (B) randomly fed blood glucose levels in WT and APPKI ^NL-G-F/NL-G-F^ mice on normal diet (ND) and HFD (n = 11–13 per group). (C) The Glucose Tolerance Test (GTT) and (D) the Insulin Tolerance Test (ITT) {12 weeks, n = 5–6 per group). (E) Fasting blood insulin levels (12 weeks, n = 4 per group). (F) Food intake and (G) rectal temperature in random-fed and 3 hours-fasted conditions (12 weeks, n = 5 per group). The blood levels of (H) total ketone body, (I) cholesterol (total, low-density lipoprotein [LDL], and high-density lipoprotein [HDL]), (J) triglyceride, and (K) free fatty acid under fasting condition (12 weeks, n = 5–6 per group). Results are presented as mean ± standard error of the mean (SEM). WT:ND vs APPKI ^NL-G-F/NL-G-F^:ND, *p < 0.05, **p < 0.01; WT:ND vs WT:HFD, †p < 0.05, ††p < 0.01; WT:ND vs APPKI ^NL-G-F/NL-G-F^:HFD, ‡p < 0.05, ‡‡p < 0.01; APPKI ^NL-G-F/NL-G-F^:ND vs APPKI ^NL-G-F/NL-G-F^:HFD, §p < 0.05, §§p < 0.01; WT:HFD vs APPKI ^NL-G-F/NL-G-F^:HFD, ¶p < 0.05, ¶¶p< 0.01.

### 3.2 HFD-induced metabolic disorders are inadequate to impair hippocampus-associated behaviors in young APPKI ^NL-G-F/NL-G-F^ mice

To evaluate the effect of HFD-induced metabolic disturbances on cognitive functions in young APPKI ^NL-G-F/NL-G-F^ mice, hippocampus-associated behavioral tests were conducted. There were no statistically significant differences in spontaneous activity or learning and memory between all groups (Figure 2A and B). Similarly, hippocampus-dependent spatial memory was comparable between ND and HFD groups, whether WT mice or APPKI ^NL-G-F/NL-G-F^ mice (Figure 2C). Furthermore, no statistically significant differences in anxiety-like behavior between all groups were observed (Figure 2D). Meanwhile, HFD-induced metabolic dysfunctions had no impact on the levels of Aβ40 and Aβ42 in the T-PER fractions of the hippocampus in young APPKI ^NL-G-F/NL-G-F^ mice, and the levels of Aβ40 consistently decreased regardless of the type of diet (Figure 2E and F). These data suggest that HFD-induced metabolic disorders are insufficient to exert a negative impact on cognitive functions and Aβ levels in APPKI ^NL-G-F/NL-G-F^ and WT mice during young adulthood.

**FIGURE 2.**
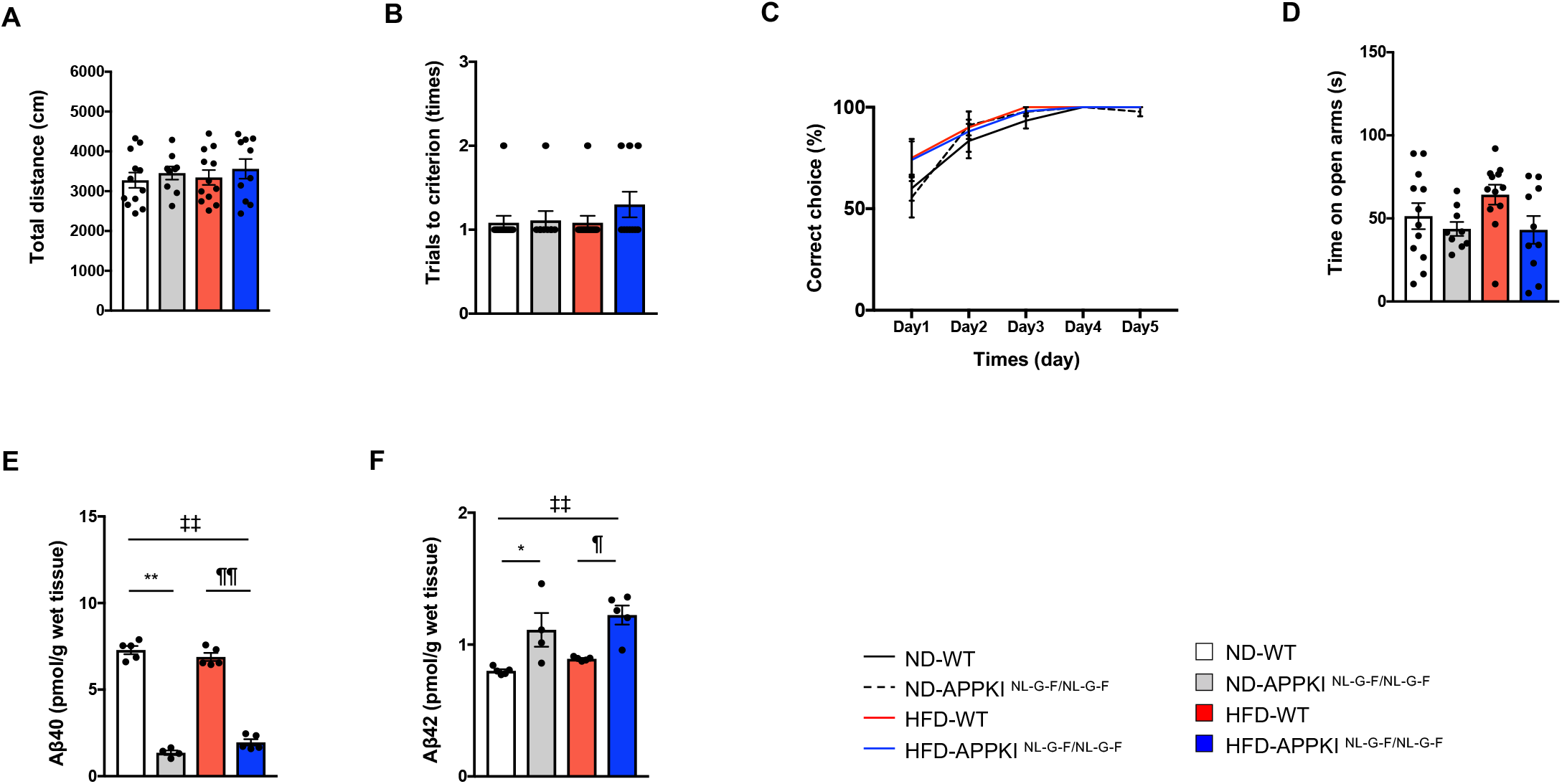
No effect of high-fat diet (HFD)-induced metabolic disorder on behaviors in young APPKI ^NL-G-F/NL-G-F^ mice. (A) Open field, (B) passive avoidance test, (C) the water T maze test, and (D) elevated plus maze in WT and APPKI ^NL-G-F/NL-G-F^ mice on normal diet (ND) and HFD (12 weeks, n = 9–12 per group). Levels of T-PER-extractable (E) Aβ40 and (F)Aβ42 in the hippocampi in the respective mouse lines (12 weeks, n = 4–5 per group). Results are presented as mean ± standard error of the mean (SEM), WT:ND vs APPKI ^NL-G-F/NL-G-F^:ND, *p < 0.05, **p < 0.01; WT:ND vs APPKI ^NL-G-F/NL-G-F^:HFD, ‡‡p < 0.01; WT:HFD vs APPKI ^NL-G-F/NL-G-F^:HFD, ¶p < 0.05, ¶¶p < 0.01.

### 3.3 HFD-young APPKI ^NL-G-F/NL-G-F^ mice display unchanged levels of specific Ser phosphorylation on hippocampal IRS1

Lastly, whether the HFD-induced metabolic disorders influence IRS1 phosphorylation in the hippocampus of young APPKI ^NL-G-F/NL-G-F^ mice on HFD was assessed. Consistent with the general change in cognitive functions and Aβ levels, no statistically significant differences in the IRS1 phosphorylation at mSer307 and mSer612 sites between both mice on ND and HFD were observed. However, these IRS1 phosphorylation levels in both types of mice on HFD were found to increase compared to both types of mice on ND (Figure 3A and B).

**FIGURE 3.**
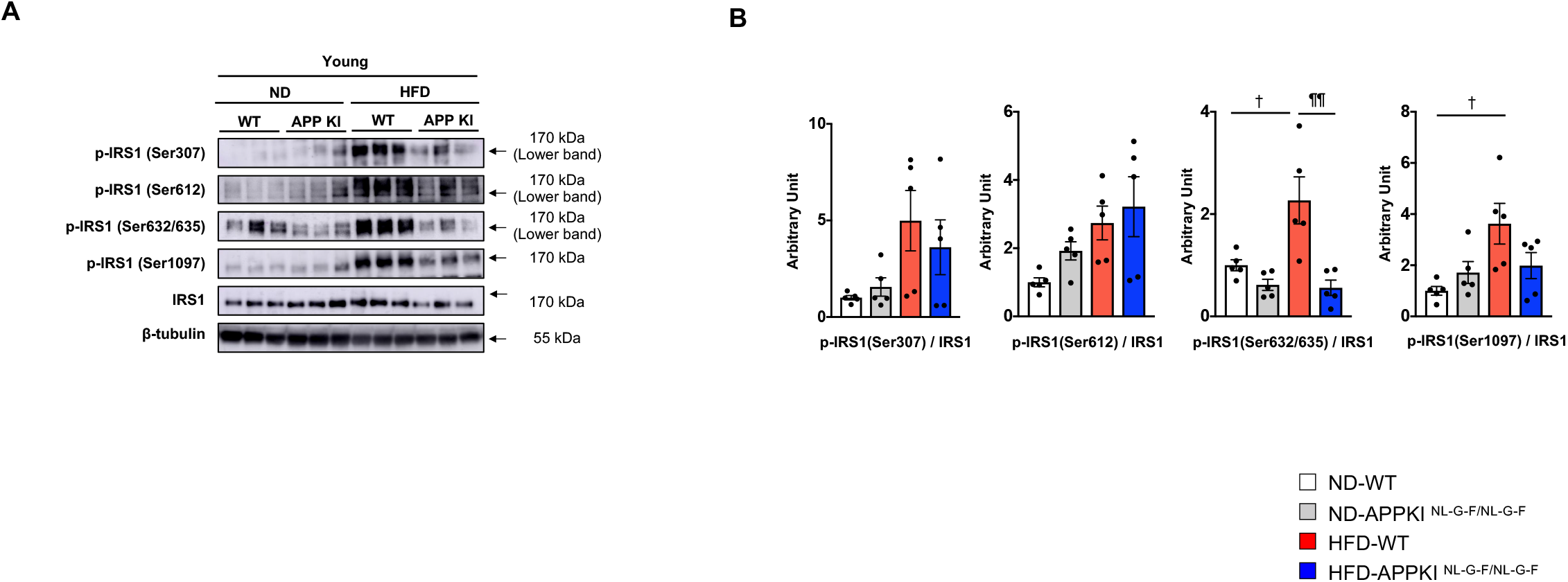
High-fat diet (HFD)-induced metabolic disturbance has no impact on the alteration of IRS1 Ser phosphorylation in the hippocampus of young APPKI ^NL-G-F/NL-G-F^ mice. Western blot analysis: (A) the phosphorylation levels of mouse Ser307 (mSer307), mSer612, mSer632/635, and mSer1097 sites on IRS1, the respective total protein levels, and β-tubulin in the hippocampi of WT and APPKI ^NL-G-F/NL-G-F^ mice on normal diet (ND) and HFD (12 weeks, n = 5 per group). (B) Graphs show relative quantification of Ser phosphorylation, respectively. An arrow indicates their respective bands in (A). The phosphorylation of their respective molecules was normalized to their respective total protein content. Results are presented as mean ± standard error of the mean (SEM), WT:ND vs WT:HFD, †p < 0.05; WT:HFD vs APPKI ^NL-G-F/NL-G-F^:HFD, s¶¶p < 0.01.

Although young WT mice responded to HFD, leading to elevated levels of IRS1phosphorylation at mSer632/635 and mSer1097 sites, these Ser sites on IRS1 in young APPKI ^NL-G-F/NL-G-F^ mice were resistant to HFD, indicating the monotonous levels of IRS1 phosphorylation (Figure 3B). Thus, HFD-induced metabolic dysfunctions do not cause increased IRS1 phosphorylation at Ser sites in the hippocampus of young APPKI ^NL-G-F/NL-G-F^ mice exhibiting normal cognitive functions.

## 4 DISCUSSION

In the present study, we have shown that both young WT and APPKI ^NL-G-F/NL-G-F^ mice on HFD display T2DM- and obesity-related typical abnormalities of nutrient metabolism, such as weight gain, impaired glucose metabolism, and hyperlipidemia, in addition to characteristic alterations of energy homeostasis in the brain, such as impaired thermoregulation and abnormal ketogenesis. Nevertheless, HFD-induced metabolic disorders are inadequate to alter the levels of Aβ and to induce cognitive deficits accompanied by the specific IRS1 modifications in both young WT and APPKI ^NL-G-F/NL-G-F^ mice.

While HFD treatment can cause T2DM- and obesity-associated metabolic disorders in WT mice regardless of mouse age and treatment duration^18-21^, several Tg-AD mouse models on ND exhibit metabolic abnormalities. For example, ND-3xTgAD mice naturally gain increased body weight and elevated blood glucose and triglyceride^12, 13^. By contrast, ND-APPKI ^NL-G-F/NL-G-F^ mice display normal metabolic parameters even after middle age similar to APPKI ^NL/N^ and APPKI ^NL-F/NL-F^ mice on ND ^7, 14, 15^. Nevertheless, young APPKI ^NL/N^ and APPKI ^NL-G-F/NL-G-F^ mice respond to HFD, showing metabolic disorders similar to young WT mice (whether young APPKI ^NL-F/NL-F^ mice respond to HFD is not reported). Given that APPKI ^NL-G-F/NL-G-F^ mice carry three mutations at mouse APP locus than none in WT mice, one mutation in APPKI ^NL/N^ mice, or two mutations in APPKI ^NL-F/NL-F^ mice, APPKI ^NL-G-F/NL-G-F^ mice are likely to have increased susceptibility to HFD, leading to more severe metabolic dysfunctions than others on HFD. However, practically there are no significant differences in metabolic alterations (i) between young WT and APPKI ^NL-G-F/NL-G-F^ mice on HFD (ii) between young or middle-aged WT and APPKI ^NL/N^ mice on HFD (iii) between middle-aged WT and APPKI ^NL-F/NL-F^mice on HFD, regardless of age, treatment stage or dietary fat ratio^14, 15^. These unexpected outcomes may be due to the low effect of HFD in young APPKI ^NL-G-F/NL-G-F^ mice, middle-aged APPKI ^NL/N^ and APPKI ^NL-F/NL-F^ mice. However, the impact of age accompanied by HFD is supposed to deteriorate further energy metabolism and neurological functions than individual effects of aging or HFD.

Thermoregulatory disorder attributed to aberrant changes in core body temperature correlates with impaired energy metabolism, which is observed even before the onset of neurodegenerative diseases, such as AD^22^. Although the rectal temperature is comparable between WT and APPKI ^NL-G-F/NL-G-F^ mice under hypernutritional states and between all groups under fasting conditions, core body temperature under ND feeding significantly decreases in APPKI ^NL-G-F/NL-G-F^ mice, suggesting that young APPKI ^NL-G-F/NL-G-F^ mice are more susceptible to impaired thermoregulation.

Ketone bodies are the lipid-derived molecules produced during fasting or on HFD as an alternative energy fuel for the brains. The ketone bodies can compensate for the deterioration of brain energy metabolism, such as lower brain glucose levels observed in neurodegenerative diseases, such as AD^23-25^. Interestingly, in young APPKI ^NL-G-F/NL-G-F^ mice, the level of blood total ketone bodies in the fasting state was constantly reduced irrespective of the diet type, which is consistent with previous studies showing that beta-hydroxybutyrate/beta-hydroxybutyric acid, is decreased in the blood and brain parenchyma of patients with AD compared to non-AD subjects^26, 27^. As persistent activation of adenosine monophosphate–activated protein kinase (AMPK) correlates with energy depletion and AD arises in the brains of young and middle-aged APPKI ^NL-G-F/NL-G-F^ mice on ND^7, 28, 29^, it is reasonable that APPKI ^NL-G-F/NL-G-F^ mice may possess deteriorating brain energy metabolism accompanied by impaired ketogenesis.

Although long-or short-term HFD-induced metabolic dysfunctions that cause Aβ-unrelated behavioral alterations in WT mice^7, 19, 21^ have no impact on hippocampus-associated behaviors in young APPKI ^NL-G-F/NL-G-F^ mice, a simple comparison in these different impacts of HFD on behaviors between WT and APPKI ^NL-G-F/NL-G-F^ mice is practically impossible. This is because behavioral alterations, above all on short-term HFD, are observed on hippocampus-independent behavioral tests or different test batteries used in APPKI mice groups^14, 15^(Figure 2A-D). Nevertheless, chronic HFD for six months or more induces hippocampus-associated behavioral abnormalities in middle-aged APPKI ^NL/NL^ and APPKI ^NL-F/NL-F^ mice regardless of alterations in Aβ levels, differences in APPKI lines, and the initiation age of HFD. However, unlike the immediate effects of HFD on nutrient metabolism, HFD for nine weeks in early life does not alter cognitive function, which could be attributed to the lack of aging effects accompanied by long-term HFD.

Additionally, given that cognitive impairment occurs regardless of alterations in Aβ levels, the correlation between T2DM and AD neuropathology remains controversial ^7, 12^-^15, 30^. Indeed, chronic HFD enhances AD neuropathology in middle-aged APPKI ^NL-F/NL-F^ mice but not in middle-aged APPKI ^NL/NL^ mice. On the other hand, HFD in early life has no impact on APPKI ^NL-G-F/NL-G-F^ mice, suggesting that HFD in early life is insufficient to influence Aβ levels in young APPKI ^NL-G-F/NL-G-F^ mice. Similarly, consistent with previous data on the behaviors and Aβ levels, HFD-induced metabolic disorders do not cause significant alterations in the levels of specific Ser phosphorylation on neural IRS1 in young APPKI ^NL-G-F/NL-G-F^ mice, whereas HFD significantly increases the neural IRS1 phosphorylation levels at two Ser residues in young WT mice despite unchanged behaviors. These data suggest that, in young APPKI ^NL-G-F/NL-G-F^ mice, Ser phosphorylation sites on neural IRS1 have reduced susceptibility to HFD compared to those in young WT mice. We have recently demonstrated that middle-aged (8 months of age) APPKI ^NL-G-F/NL-G-F^ mice display cognitive deficits with persistent IRS1 phosphorylation at specific Ser sites, elevated Aβ42 levels, and constant activation of hippocampal AMPK, also observed in young APPKI ^NL-G-F/NL-G-F^ mice ^7^.

This indicates that consistent deterioration in brain energy metabolism triggers cognitive dysfunctions in APPKI ^NL-G-F/NL-G-F^ mice. The present study indicates that young APPKI ^NL-G-F/NL-G-F^ mice on HFD exhibit impaired ketogenesis associated with defective brain energy metabolism accompanied by abnormal glucose metabolism, dyslipidemia, and thermoregulatory disorder. However, normal cognitive function and unchanged levels of specific Ser phosphorylation on neural IRS1 and of Aβ suggest that HFD in early life is insufficient to cause neurological alterations, which are likely to be triggered by the aging effect with chronic HFD. Further studies need to be conducted to determine the impact of long-term HFD on nutrient metabolism and neurological phenotypes in APPKI ^NL-G-F/NL-G-F^ mice after middle age.

## ACKNOWLEDGMENTS

We thank Kohei Tomita and Dr. Noboru Ogiso for animal care and Dr. Yudai Shibayama for preparing and helping behavioral studies. This work is supported by grants from the Ministry of Education, Culture, Sports, Science and Technology (20H04137) (A.T.), Research supports from DAIICHI SANKYO COMPANY, LIMITED (A.T.) and SHIONOGI & CO., LTD (A.T.), and the Research Funding for Longevity Science from the National Center for Geriatrics and Gerontology, Japan (A.T.).

## AUTHOR CONTRIBUTIONS

W.W., D.T., M.M., and C.K. researched data. W.W. and D.T. analyzed data. W.W. and D.T. wrote the manuscript. T.S. and T.C.S. provided the APPKI mice and interpreted the data. A.T. designed experiments, analyzed data, and wrote and edited the manuscript. All authors read and approved the final manuscript.

## DATA AVAILABILITY STATEMENT

The data that support the findings of this study are available in Supporting information.

## ANIMAL STUDIES

Animal experiments were conducted in compliance with the guidelines and approval of the ethics committee in Animal Care and Use of the National Center for Geriatrics and Gerontology in Japan.

## REFERENCES

1. Biessels GJ, Koffeman A, Scheltens P. Diabetes and cognitive impairment. Clinical diagnosis and brain imaging in patients attending a memory clinic. J Neurol 2006; 253(4): 477-82.SSSS

2. Ninomiya T. Epidemiological Evidence of the Relationship Between Diabetes and Dementia. Adv Exp Med Biol 2019; 1128: 13–25.

3. Copps KD, White MF. Regulation of insulin sensitivity by serine/threonine phosphorylation of insulin receptor substrate proteins IRS1 and IRS2. Diabetologia 2012; 55(10): 2565–2582.

4. Moloney AM, Griffin RJ, Timmons S, O’Connor R, Ravid R, O’Neill C. Defects in IGF-1 receptor, insulin receptor and IRS-1/2 in Alzheimer’s disease indicate possible resistance to IGF-1 and insulin signalling. Neurobiology of aging 2010; 31(2): 224–43.

5. Talbot K, Wang HY, Kazi H, Han LY, Bakshi KP, Stucky A et al. Demonstrated brain insulin resistance in Alzheimer’s disease patients is associated with IGF-1 resistance, IRS-1 dysregulation, and cognitive decline. The Journal of clinical investigation 2012; 122(4): 1316–38.

6. Saito T, Matsuba Y, Mihira N, Takano J, Nilsson P, Itohara S et al. Single App knock-in mouse models of Alzheimer’s disease. Nature neuroscience 2014; 17(5): 661–3.

7. Wang W, Tanokashira D, Fukui Y, Maruyama M, Kuroiwa C, Saito T et al. Serine Phosphorylation of IRS1 Correlates with Aβ-Unrelated Memory Deficits and Elevation in Aβ Level Prior to the Onset of Memory Decline in AD. Nutrients 2019; 11(8).

8. Beeri MS, Silverman JM, Davis KL, Marin D, Grossman HZ, Schmeidler J et al. Type 2 diabetes is negatively associated with Alzheimer’s disease neuropathology. The journals of gerontology. Series A, Biological sciences and medical sciences 2005; 60(4): 471–5.

9. Matsuzaki T, Sasaki K, Tanizaki Y, Hata J, Fujimi K, Matsui Y et al. Insulin resistance is associated with the pathology of Alzheimer disease: the Hisayama study. Neurology 2010; 75(9): 764–70.

10. Arnold SE, Arvanitakis Z, Macauley-Rambach SL, Koenig AM, Wang HY, Ahima RS et al. Brain insulin resistance in type 2 diabetes and Alzheimer disease: concepts and conundrums. Nature reviews. Neurology 2018; 14(3): 168–181.

11. Ho L, Qin W, Pompl PN, Xiang Z, Wang J, Zhao Z et al. Diet-induced insulin resistance promotes amyloidosis in a transgenic mouse model of Alzheimer’s disease. FASEB journal: official publication of the Federation of American Societies for Experimental Biology 2004; 18(7): 902–4.

12. Vandal M, White PJ, Tremblay C, St-Amour I, Chevrier G, Emond V et al. Insulin reverses the high-fat diet-induced increase in brain Aβ and improves memory in an animal model of Alzheimer disease. Diabetes 2014; 63(12): 4291–301.

13. Knight EM, Martins IV, Gümüsgöz S, Allan SM, Lawrence CB. High-fat diet-induced memory impairment in triple-transgenic Alzheimer’s disease (3xTgAD) mice is independent of changes in amyloid and tau pathology. Neurobiology of aging 2014; 35(8): 1821–32.

14. Salas IH, Weerasekera A, Ahmed T, Callaerts-Vegh Z, Himmelreich U, D’Hooge R et al. High fat diet treatment impairs hippocampal long-term potentiation without alterations of the core neuropathological features of Alzheimer disease. Neurobiol Dis 2018; 113: 82–96.

15. Mazzei G, Ikegami R, Abolhassani N, Haruyama N, Sakumi K, Saito T et al. A high-fat diet exacerbates the Alzheimer’s disease pathology in the hippocampus of the App(NL-F/NL-F) knock-in mouse model. Aging cell 2021; 20(8): e13429.

16. Petrov D, Pedrós I, Artiach G, Sureda FX, Barroso E, Pallàs M et al. High-fat diet-induced deregulation of hippocampal insulin signaling and mitochondrial homeostasis deficiences contribute to Alzheimer disease pathology in rodents. Biochimica et biophysica acta 2015; 1852(9): 1687–99.

17. Tanokashira D, Wang W, Maruyama M, Kuroiwa C, White MF, Taguchi A. Irs2 deficiency alters hippocampus-associated behaviors during young adulthood. Biochemical and biophysical research communications 2021; 559: 148–154.

18. Tanokashira D, Kurata E, Fukuokaya W, Kawabe K, Kashiwada M, Takeuchi H et al. Metformin treatment ameliorates diabetes-associated decline in hippocampal neurogenesis and memory via phosphorylation of insulin receptor substrate 1. FEBS Open Bio 2018; 8(7): 1104–1118.

19. Gainey SJ, Kwakwa KA, Bray JK, Pillote MM, Tir VL, Towers AE et al. Short-Term High-Fat Diet (HFD) Induced Anxiety-Like Behaviors and Cognitive Impairment Are Improved with Treatment by Glyburide. Front Behav Neurosci 2016; 10: 156.

20. Johnson LA, Zuloaga KL, Kugelman TL, Mader KS, Morré JT, Zuloaga DG et al. Amelioration of Metabolic Syndrome-Associated Cognitive Impairments in Mice via a Reduction in Dietary Fat Content or Infusion of Non-Diabetic Plasma. EBioMedicine 2016; 3: 26–42.

21. McLean FH, Grant C, Morris AC, Horgan GW, Polanski AJ, Allan K et al. Rapid and reversible impairment of episodic memory by a high-fat diet in mice. Scientific reports 2018; 8(1): 11976.

22. Cheshire WP, Jr. Thermoregulatory disorders and illness related to heat and cold stress. Auton Neurosci 2016; 196: 91–104.

23. Jensen NJ, Wodschow HZ, Nilsson M, Rungby J. Effects of Ketone Bodies on Brain Metabolism and Function in Neurodegenerative Diseases. Int J Mol Sci 2020; 21(22).

24. Cunnane SC, Courchesne-Loyer A, St-Pierre V, Vandenberghe C, Pierotti T, Fortier M et al. Can ketones compensate for deteriorating brain glucose uptake during aging? Implications for the risk and treatment of Alzheimer’s disease. Ann N Y Acad Sci 2016; 1367(1): 12–20.

25. García-Rodríguez D, Giménez-Cassina A. Ketone Bodies in the Brain Beyond Fuel Metabolism: From Excitability to Gene Expression and Cell Signaling. Front Mol Neurosci 2021; 14: 732120.

26. Shippy DC, Wilhelm C, Viharkumar PA, Raife TJ, Ulland TK. β-Hydroxybutyrate inhibits inflammasome activation to attenuate Alzheimer’s disease pathology. J Neuroinflammation 2020; 17(1): 280.

27. Newman JC, Verdin E. β-Hydroxybutyrate: A Signaling Metabolite. Annu Rev Nutr 2017; 37: 51–76.

28. Liu YJ, Chern Y. AMPK-mediated regulation of neuronal metabolism and function in brain diseases. J Neurogenet 2015; 29(2-3): 50–8.

29. Wang X, Zimmermann HR, Ma T. Therapeutic Potential of AMP-Activated Protein Kinase in Alzheimer’s Disease. Journal of Alzheimer’s disease: JAD 2019; 68(1): 33–38.

30. Rabinovici GD, Furst AJ, Alkalay A, Racine CA, O’Neil JP, Janabi M et al. Increased metabolic vulnerability in early-onset Alzheimer’s disease is not related to amyloid burden. Brain 2010; 133(Pt 2): 512–28.

