## Supplementary figures and images for "Young APPKI ^NL-G-F/NL-G-F^ mice display high-fat diet-induced metabolic disturbances and specific disorders associated with brain energy homeostasis"

### Supplementary Figure 1

#### SUPPLEMENTAL FILE

##### Supplementary Figure 1 for Figure 3A

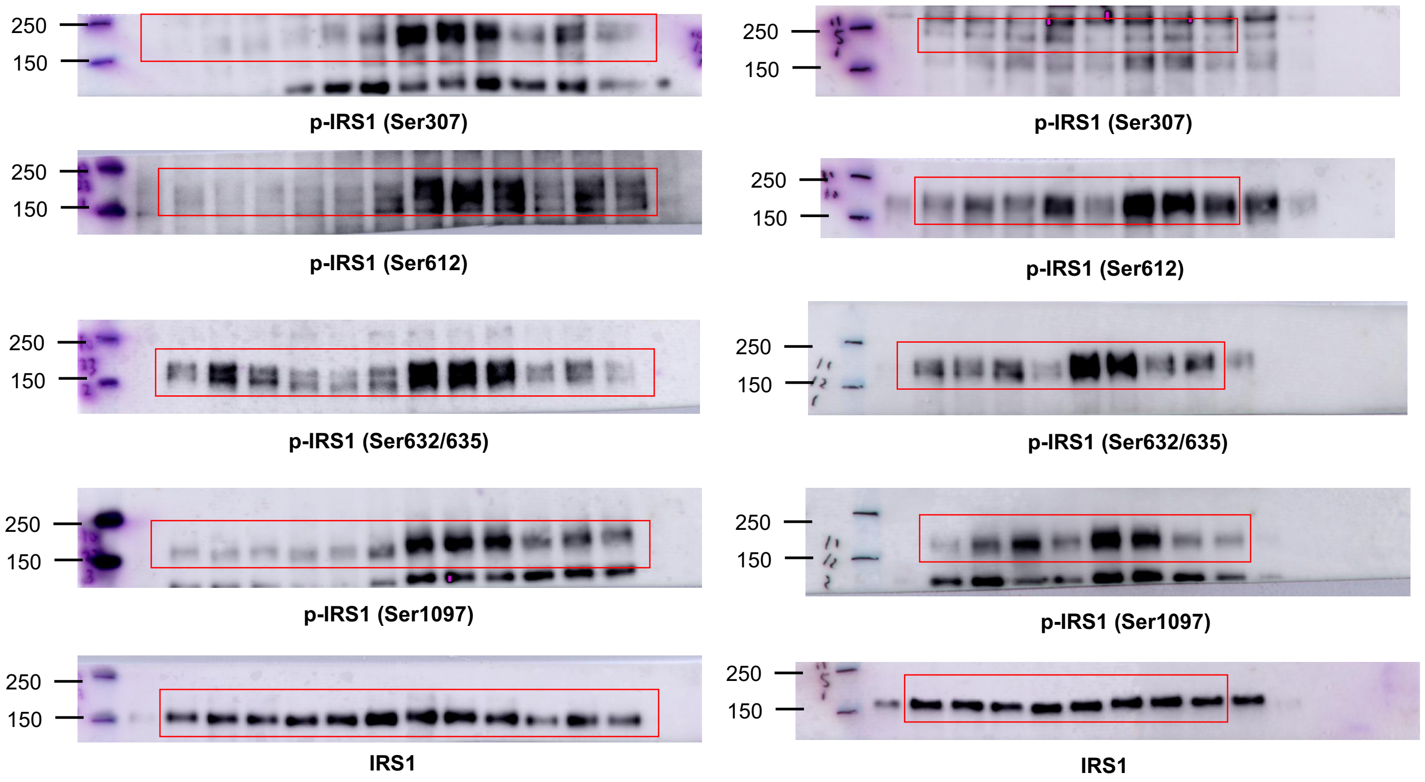
